# Modulation of mitochondrial Hsp90 (TRAP1) ATPase activity by calcium and magnesium

**DOI:** 10.1101/300038

**Authors:** Daniel Elnatan, David A. Agard

**Affiliations:** Howard Hughes Medical Institute and the Department of Biochemistry & Biophysics University of California, San Francisco San Francisco, CA 94143

## Abstract

The Hsp90 protein family are ATP-dependent molecular chaperones that maintain protein homeostasis and regulate many essential cellular processes. Higher eukaryotic cells have organelle-specific Hsp90 paralogs that are adapted to each unique sub-cellular environment. The mitochondrial Hsp90, TRAP1, supports the folding and activity of electron transport components and is increasingly being appreciated as a critical player in mitochondrial signaling. It is well known that calcium plays an important regulatory role in mitochondria and can even accumulate to much higher concentrations than in the cytoplasm. Surprisingly, we find that calcium can replace the requirement for magnesium to support TRAP1 ATPase activity. Using anomalous x-ray diffraction, we reveal a novel calcium-binding site within the TRAP1 nucleotide-binding pocket located near the ATP α-phosphate and completely distinct from the magnesium site adjacent to the β and γ-phosphates. In the presence of magnesium, ATP hydrolysis by TRAP1, as with other Hsp90s, is non-cooperative, whereas calcium binding results in cooperative ATP hydrolysis by the two protomers within the Hsp90 dimer. The structural data suggest a mechanism for the cooperative behavior. Owing to the cooperativity, at high ATP concentrations, ATPase activity is higher with calcium, whereas the converse is true at low ATP concentrations. Integrating these observations, we propose a model where the divalent cations choice can control switching between non-cooperative and cooperative TRAP1 ATPase mechanisms in response ATP concentrations. This may facilitate coordination between cellular energetics, mitochondrial signaling, and protein homeostasis via alterations in the TRAP1 ATP-driven cycle.

## Introduction

Enzymatic studies of ATPases generally include magnesium as the physiological co-factor required for both nucleotide binding as well as catalysis. Except in specific cases, other divalent ions are generally not considered as even calcium concentrations are kept quite low (nanomolar) in the resting cytoplasm. By contrast with the cytoplasm, sub-cellular organelles can have vastly different ionic environments. It is now well appreciated that mitochondria serve as a major cellular calcium reservoir in non-muscle cells. Calcium-phosphate granules form inside the mitochondrial matrix facilitating calcium storage and buffering, and if completely dissolved, would reach molar concentrations^1^. Many studies show that calcium levels within mitochondria are tightly linked to both mitochondrial and cellular signaling pathways, including cell death via the mitochondrial permeability transition pore^2^. Unsurprisingly, several mitochondrial enzymes are regulated by calcium ions^3^. In particular, mitochondrial dehydrogenases involved in ATP production are stimulated by calcium ions, suggesting a direct link to cellular energetics. Given the prevalence of calcium in the mitochondrial matrix, it is likely that calcium ions play important roles in regulating other mitochondrial proteins. In this study, we look at how a mitochondrial homolog of the molecular chaperone Hsp90 (TRAP1) may be regulated by calcium binding via a novel mechanism.

Hsp90 molecular chaperones facilitate the folding and activation of ∼10% of cellular proteins^4^. Increasingly, it is appreciated that Hsp90s can support function throughout a protein’s lifetime, not just during initial synthesis or in response to cellular stress. While Hsp90s are well conserved from bacteria to mammals, distinct orthologs can be found within organelles: Grp94 in the endoplasmic reticulum, TRAP1 in mitochondria, and Hsp90.5 in chloroplasts. Beyond their role in supporting individual “client” substrate proteins, cytosolic Hsp90s are critically important for maintaining protein homeostasis and modulating certain pro-apoptotic kinases.

All Hsp90s are homodimers with each protomer composed of 3 globular domains: an ATP-binding N-terminal Domain (NTD), a middle domain (MD), and C-terminal dimerizing domain (CTD). Differences serve to adapt the Hsp90s for their various cellular roles, interactions with co-chaperones (cytoplasm only) and localizations. Hsp90s go through complex conformational rearrangements (domain rotations and open-closed transitions) throughout their ATP hydrolysis cycle^5^ and this ATPase activity is absolutely essential for its function in eukaryotic cells^6,7^. Despite being structurally conserved, Hsp90 homologs differ in their ATPase kinetics^8-10^, likely reflecting a distinct yet precisely tuned conformational equilibrium between the open and closed state for each species^11^.

While the *in*-*vivo* roles of cytosolic Hsp90s have been the most studied owing to their importance for disease and evolution^12^, recently attention is being focused on the mitochondrial isoform, TRAP1. TRAP1 has been implicated in regulating mitochondrial energetics^13^, the opening of the mitochondrial transition pore^14^ and kinase signaling^15^ suggesting linkages between mitochondrial protein homeostasis, energetics and broader aspects of cell function. Also relevant, have been clear connections to neurodegeneration^16,17^ and cancer^18^.

TRAP1 structural studies have revealed the presence of a remarkably asymmetric closed state in the presence of non-hydrolyzable ATP analogs^19^ that leads to a two-step sequential ATP hydrolysis mechanism, with the first ATP leading to a flip in asymmetry, and the second leading to re-opening (Figure 4)^20^. As the primary client binding site maps to the region of maximal asymmetry, this suggests a strong coupling between the asymmetry flip and client remodeling. Another unique feature of TRAP1 is an N-terminal extension that renders its open-closed equilibrium particularly sensitive to temperature variations, possibly connecting its folding and signaling role to mitochondrial thermogenesis^21^.

While ATP is required to stabilize the closed N-terminally dimerized state in all Hsp90s, uniquely for TRAP1 closure (but not hydrolysis) will occur in the absence of any divalent cation co-factor^20,21^. Although the mechanism and purpose of this phenomenon are unknown, there have been reports of free ATP (not complexed to Mg^2+^) existing in the mitochondrial matrix^22^.

To obtain a better understanding how divalent cations may regulate TRAP1, experimental systems with purified protein can bypass the complexity of cell-based models; providing useful mechanistic insights that may be important in the cellular context. To this end, we investigated how the ATPase activity of TRAP1 is regulated by two divalent cations: magnesium or calcium. We found that unlike most other Hsp90 homologs, the TRAP1 ATPase can efficiently utilize calcium alone to support hydrolysis, and that the relative preference for magnesium or calcium depends upon the concentration of free ATP. Moreover, calcium induced a profound change in the cooperativity of ATP hydrolysis, indicating a change in the underlying mechanism. Crystallographic analysis revealed a calcium binding site within the NTD distinct from that used by magnesium. These observations lead to a model of how cation binding within TRAP1 can influence the TRAP1 catalytic cycle in an ATP-level dependent manner.

## Results

### TRAP1 has unique enzyme kinetics depending on magnesium or calcium

Given the role of mitochondria as a cellular calcium reservoir, we decided to test whether calcium might affect ATP hydrolysis by human TRAP1. ATP hydrolysis by Hsp90 happens in the closed state when the NTDs are dimerized. This closed state can be stabilized by non-hydrolysable ATP analogs, or in the case of TRAP1, simply by omitting magnesium. Under these conditions, the rate of ATP turnover is much slower than the usually rate-limiting dimer closure, resulting in a build-up of the closed state^20,21^. We have previously taken advantage of this phenomenon to decouple dimer closure from ATP hydrolysis, which can be initiated by addition of Mg^2+^. The closed dimer reopens once ATP hydrolysis resumes, and this can be readily monitored by a FRET assay (Figure 1A). Since the steady-state assay only reads out the slow step of dimer closure, we decided to test whether calcium alone can trigger ATP hydrolysis from the closed state. As previously observed, addition of magnesium causes loss of FRET signal due to hydrolysis-triggered dimer re-opening. The addition of calcium can also cause dimer opening (Figure 1B), and with a ∼10-fold faster kinetics than magnesium. This suggests that in the absence of magnesium, calcium can fully support ATP hydrolysis by TRAP1 once it is closed.

**Figure 1.**
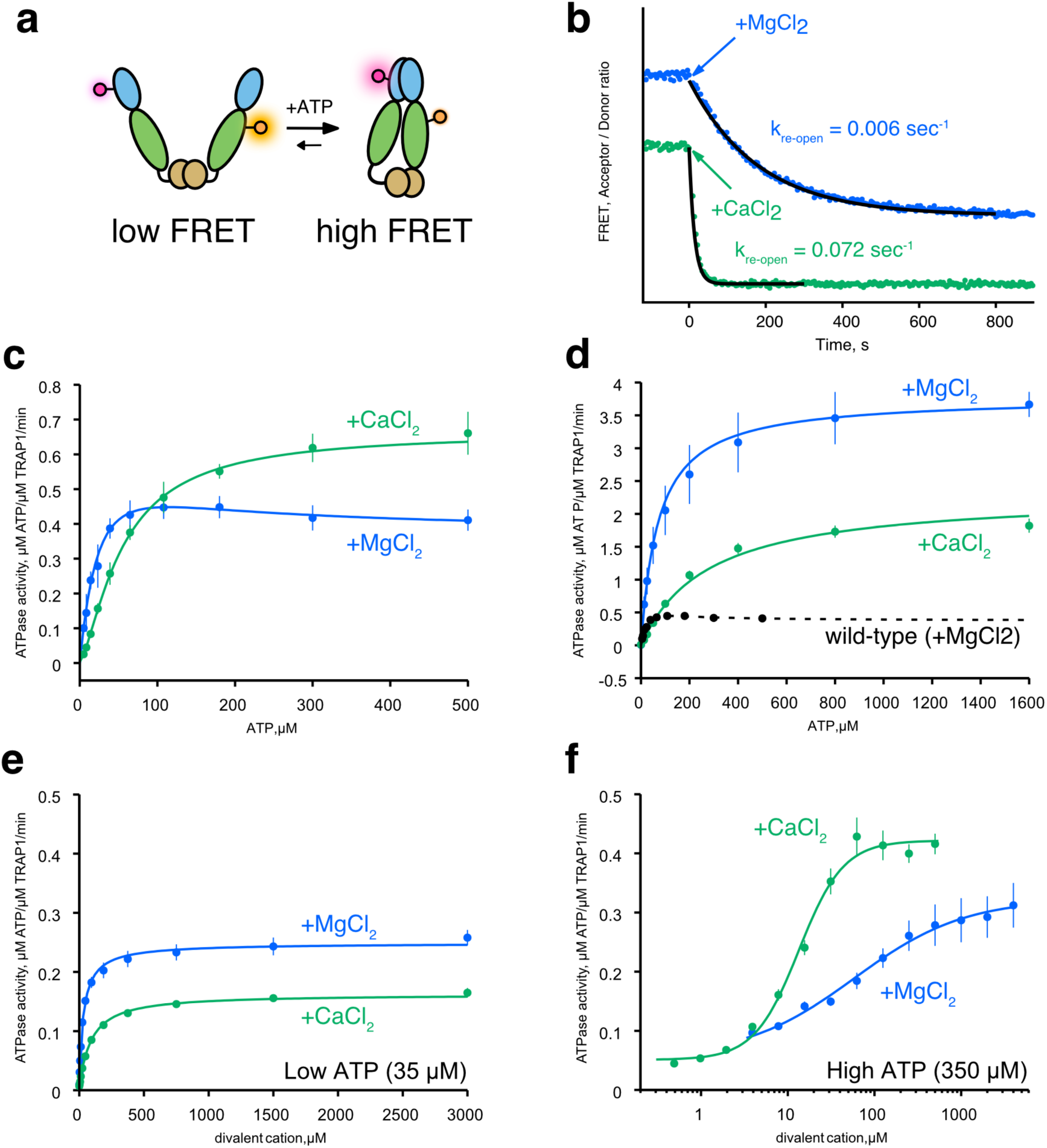
Calcium supports ATP hydrolysis by TRAP1. **a**) Cartoon diagram of the FRET assay with TRAP1 to monitor open (low FRET) and closed state (high FRET). Dye molecules (glowing circles) are covalently attached at the middle domain (green ellipse) and the NTD (blue ellipse). **b**) A representative FRET re-opening assay starting from the closed state (high FRET). Reaction was initiated by addition of either MgCl_2_ (blue) or CaCl_2_ (green) at t=0. Time-dependent decrease of FRET signal indicate dimer opening. FRET is quantified as the ratio of Acceptor/Donor intensity. The kinetic trace has been vertically offset for clarity. Black lines are exponential fits to the re-opening kinetics with indicated rate constants, k_re-open_. Measurements were repeated at least three times. **c**) ATPase activity of hTRAP1 in presence of excess MgCl_2_(blue) or CaCl_2_(green) as a function of ATP concentration. **d**) ATPase activity of wild-type:D158N heterodimeric mixture in presence of MgCl_2_ or CaCl_2_ as a function of ATP concentration. Wild-type ATPase data with MgCl2 (black circles) and fit (dashed line) is shown for comparison. **e**) ATPase activity of hTRAP1 at low ATP concentration (35 μM) as a function of divalent cation concentrations (MgCl_2_ in blue; CaCl_2_ in green). **f**) ATPase activity of hTRAP1 at high ATP concentration (350 μM) as a function of divalent cation concentrations (MgCl_2_ in blue; CaCl_2_ in green). The error bars from Figure 1D-F are standard deviations calculated from three technical replicates.

Next, we assessed whether calcium alone can support both closure and ATP hydrolysis. As previously observed^9^, ATPase activity in the presence of magnesium is non-cooperative and reaches a plateau with as little as ∼50 μM ATP. By contrast, ATPase activity using calcium follows the Hill-equation with n=1.4 and an ATPase rate that is 1.6-fold faster than in magnesium at saturating ATP (Figure 1C, see Table 1 for fitted parameters). While fitting the data, we noted that the Michaelis-Menten equation is a poor fit to the magnesium ATPase activity (Supplemental Figure 1). This behavior also appears to be conserved in zebrafish TRAP1 (zTRAP1) (Supplemental Figure 2). The “bump” on the curve between 0 to ∼200 μM ATP can be explained by a model in which the ATPase activity of the singly bound ATP state is greater than when ATPs are bound by both protomers. We confirmed this by measuring the ATPase activity of a heterodimeric mixture of hTRAP1 in which one ATP binding site was inactivated via a D158N mutation. In the presence of magnesium, the ATPase activity curve of the heterodimer no longer has the “bump” and follows standard single-site Michaelis-Menten kinetics (Figure 1D, blue line). The apparent affinity to ATP (*K_0.5 ATP_*, defined as the concentration at half-max of activity) measured in this heterodimer (Table 3) is close to that estimated by the wild-type experiment (Table 1) (58 μM vs 73 μM). As predicted, this half-occupied ATP heterodimeric mixture has a dramatically higher ATPase activity (3.8 min^−1^) than the fully-occupied wild-type dimer (0.37 min^−1^) (Figure 1D, Table 3 and 1). This is consistent with previous heterodimer experiments where a related point mutant, which should also abolish ATP binding (D158A), appeared to stimulate ATPase activity^23^. The mechanism underlying this non-trivial rate enhancement is currently unknown, and it highlights yet another unique aspect of TRAP1 regulation by nucleotide. In contrast to the wild-type enzyme, the heterodimer ATPase activity in the presence of calcium is significantly reduced (0.6-fold), and consequently calcium is no longer preferred. To further examine whether the differential effects of magnesium versus calcium are a unique feature of TRAP1, ATPase activities of several Hsp90 orthologs were also measured. Beyond the general trend that ATPase activity is lower when calcium is used, none of the tested homologs exhibited the distinct ATP concentration-dependent activity as seen in TRAP1 (Supplemental Figure 3).

**Table 1.** Fitted parameters for enzyme kinetics data as a function of ATP concentration with excess divalent cations. The hill-coefficient (n_ATP_) and apparent affinity (K_0.5 ATP_) are for ATP in excess divalent cations. K_singleATP_ and K_doubleATP_ are the apparent affinity constants used in a two-population ATPase model for one- and two-ATP bound dimer.

|  | 37°C |  | 30°C |  |
| --- | --- | --- | --- | --- |
|  | hTRAP1, MgCl<sub>2</sub> | hTRAP1, CaCl<sub>2</sub> | zTRAP1, MgCl<sub>2</sub> | zTRAP1, CaCl<sub>2</sub> |
| **V<sub>max</sub>** | - | 0.67 ± 0.01 | - | 0.82 ± 0.01 |
| **V<sub>singleATP</sub>** | 0.68 ± 0.05 | - | 1.50 ± 0.24 | - |
| **V<sub>doubleATP</sub>** | 0.37 ± 0.02 | - | 0.56 ± 0.03 | - |
| **K<sub>0.5 ATP</sub>** | - | 56.1 ± 2.2 | - | 66.2 ± 2.5 |
| **K<sub>singleATP</sub>, μM** | 58.0 ± 7.8 | - | 126.3 ± 30.5 | - |
| **K<sub>doubleATP</sub>, μM** | 38.0 ± 19.9 | - | 25.2 ± 12.7 | - |
| **n<sub>ATP</sub>** | - | 1.4 ± 0.0 | - | 1.8 ± 0.1 |

### ATP-site occupancy biases preference for magnesium versus calcium

To better understand the inter-relationship between ATPase activity and divalent cation concentrations, we examined the effects of cation concentration under high and low ATP concentrations (hTRAP1 in Figure 1E and 1F; zTRAP1 in Supplemental Figure 4). The apparent affinity (K_0.5 ion_) and occupancy (n_ion_) of the divalent cations can be approximated by fitting the Hill-equation to initial rates from titrations at fixed ATP concentrations. At low ATP concentrations, hTRAP1 has a 2.8-fold tighter apparent affinity for magnesium than calcium in (Mg^2+^, 33 μM vs. Ca^2+^, 92 μM) while zTRAP1 has a 1.7-fold higher affinity (Mg^2+^, 39 μM vs Ca^2+^, 66 μM), and also ∼1.5-fold higher activity in both species (Table 2, low ATP). At low ATP concentrations, the ion-binding Hill-coefficient is near 1.0 for both magnesium and calcium, suggesting that the cations preferentially bind to the protomer having one ATP bound. By contrast at high ATP concentrations, calcium binds ∼4.5-fold more tightly than magnesium in hTRAP1 (Mg^2+^, 59 μM vs Ca^2+^, 13 μM) and ∼7-fold in zTRAP1 (Mg^2+^, 260 μM vs Ca^2+^, 37 μM; see also Table 2, high ATP). The observed differences in apparent affinities between species most likely reflect intrinsic properties of each protein. At high ATP concentrations, *n* remains close to 1.0 for magnesium while indicating very significant cooperativity (n=1.7) for calcium.

**Table 2.** Fitted parameters for divalent cation titrations in low vs high ATP concentrations for zebrafish and human TRAP1. The Hill-coefficient (n) and apparent affinity (K_0.5 ion_) here are for ion-binding at indicated ATP concentrations.

|  | Low ATP (50 μM) |  | High ATP (400 μM) |  |
| --- | --- | --- | --- | --- |
|  | zTRAP1, MgCl<sub>2</sub> | zTRAP1, CaCl<sub>2</sub> | zTRAP1, MgCl<sub>2</sub> | zTRAP1, CaCl<sub>2</sub> |
| **V<sub>max</sub>** | 0.47 ± 0.09 | 0.29 ± 0.01 | 0.49 ± 0.02 | 0.62 ± 0.02 |
| **K<sub>0.5 ion</sub>, μM** | 38.6 ± 15.0 | 66.1 ± 5.0 | 258.9 ± 15.1 | 37.1 ± 1.6 |
| **n<sub>ion</sub>** | 0.9 ± 0.1 | 1.2 ± 0.1 | 1.6 ± 0.1 | 1.9 ± 0.1 |
|  | Low ATP (35 μM) |  | High ATP (350 μM) |  |
|  | hTRAP1, MgCl<sub>2</sub> | hTRAP1, CaCl<sub>2</sub> | hTRAP1, MgCl<sub>2</sub> | hTRAP1, CaCl<sub>2</sub> |
| **V<sub>max</sub>** | 0.24 ± 0.00 | 0.16 ± 0.00 | 0.27 ± 0.01 | 0.37 ± 0.02 |
| **K<sub>0.5 ion</sub>, μM** | 32.9 ± 1.8 | 91.9 ± 3.7 | 58.8 ± 10.5 | 13.4 ± 1.2 |
| **n<sub>ion</sub>** | 1.0 ± 0.0 | 1.1 ± 0.0 | 0.7 ± 0.1 | 1.7 ± 0.2 |

**Table 3.** Fitted parameters for ATP titration kinetics in human TRAP1 wild-type:D158N heterodimer mixture. The K_0.5_ here reports on the apparent affinity constant for ATP in the Michaelis-Menten equation. ^∗^The unit for V_max_ is expressed in μM ATP·min^−1^ / μM ATPase, and the unit for K_0.5_ is expressed in μM. The standard error values from fitting are shown for each parameter.

|  | **MgCl<sub>2</sub>** | **CaCl<sub>2</sub>** |
| --- | --- | --- |
| **V<sub>max</sub>** | 3.77 ± 0.06 | 2.31 ± 0.15 |
| **K<sub>0.5 ATP</sub>, μM** | 72.5 ± 4.4 | 273.2 ± 39.2 |

### Calcium binds to the N-terminal Domain (NTD) of TRAP1

Crystal structures of the closed state of TRAP1 clearly reveal a single magnesium binding site on each protomer, coordinating the β- and γ-phosphates of ATP as is commonly observed in many ATPases. The unitary Hill-coefficient for both ATP and magnesium likely reflects the obligatory sequential (non-cooperative) nature of ATP hydrolysis previously observed ^20^. Identifying where calcium binds would greatly facilitate understanding how calcium imparts an apparent cooperativity in TRAP1 ATPase activity, which has never been observed in other Hsp90 homologs.

As TRAP1 does not contain any apparent calcium-binding motifs, one possibility is that calcium binds at the same site used by magnesium, for example as found in several other ATPases (Hsp70/DnaK/actin family) ^24,25^. To determine where calcium binds in the closed state and any consequences on the structure, we replaced the magnesium in our standard zTRAP1 crystallization conditions ^19^ with calcium. The structure was solved to a resolution of 2.6 Å using molecular replacement, and reveals a virtually identical asymmetric closed state^19^ (Figure 2A). Despite the similarity in overall structure, no density coordinating the β- and γ-phosphates, where magnesium would be, was observed (Supplemental Figure 5a-b). To unambiguously determine where calcium binds, we exploited the anomalous X-ray scattering signal from calcium and collected two datasets: one at 7450 eV (1.664 Å) and 11111 eV (1.116 Å). Although the anomalous difference at these energies is considerably weaker than what would be required for experimental phasing significant peaks can be reliably observed for a well-determined structure using ANODE^26^. Using phases from the solved structure without the contribution of calcium atoms, the resulting anomalous difference density map reveals one strong peak within the nucleotide pocket of each protomer, located just above the α-phosphate (Figure 2A, inset). The density is tightly coordinated by six oxygen atoms in an octahedral arrangement near the “lid” of the NTD (4 backbone carbonyl oxygens, a tyrosine hydroxyl, and the α-phosphate) (Figure 2B). As a control, anomalous data was collected from crystals lacking both calcium and magnesium prepared for another study^20^. In these crystals excess EDTA was used and the dimer was closed with ATP instead of AMPPNP. No significant anomalous peaks were detected near the α-phosphate (Supplemental Figure 5c), again providing strong support that the peak observed in Figure 2A is indeed from a calcium atom.

**Figure 2.**
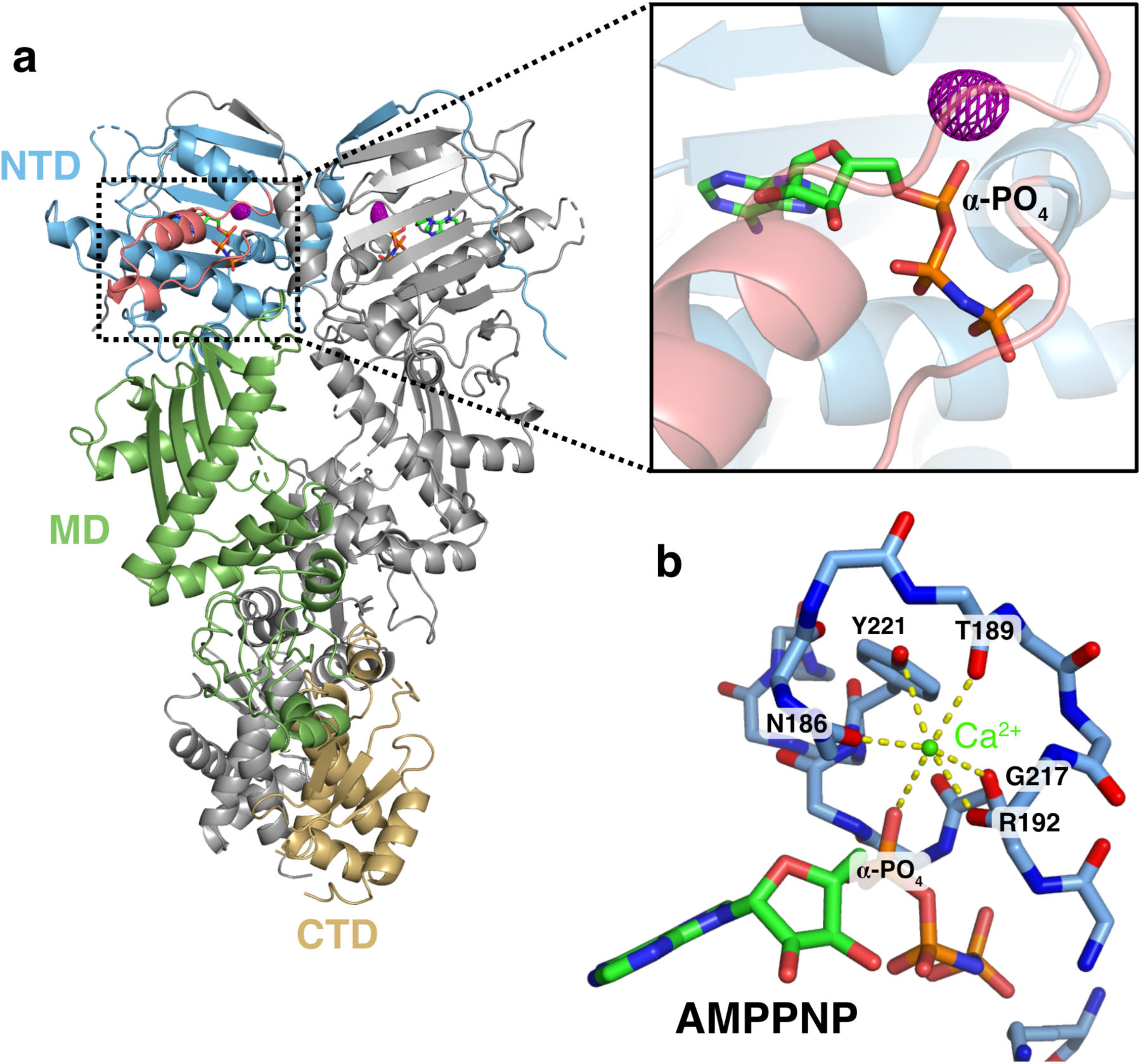
Calcium binds to the N-terminal domain of TRAP1. **a**) Full-length structure of zTRAP1 bound to AMPPNP and Ca^2+^. One protomer is colored in gray and the other is colored by domains: N-terminal domain (NTD, blue), Middle Domain (MD, green), and C-terminal Domain (CTD, gold). The ATP-lid region of the NTD is colored in pink (black rectangular inset). The anomalous electron density map (displayed at 4.5σ contour level) for calcium is shown in purple mesh. **b**) Coordination of calcium (green sphere) by residues in the ATP-lid and the α-phosphate of the AMPPNP. Yellow dashed lines depict contacts from oxygens to calcium atom. All of the oxygens were from backbone carbonyl of labeled residues with the exception of one hydroxyl oxygen from Tyr221.

## Discussion

Enzymatic studies of Hsp90 have almost exclusively used magnesium as a co-factor. Following up on an unexpected observation that calcium alone was sufficient to support TRAP1 ATPase activity, here we show that TRAP1 has a higher ATPase activity in the presence of calcium than in magnesium. This contrasts sharply with other Hsp90 orthologs, which are significantly less active with calcium. Strikingly, there are also significant differences in ATPase behavior as a function of ATP and cation concentration, with calcium inducing a cooperative dependence on ATP. Furthermore, at high ATP levels, activity cooperatively depends on calcium concentration, whereas magnesium acts non-cooperatively. Consequently, calcium can differentially regulate TRAP1 activity in an ATP-concentration dependent manner. Together this reveals that these ions must act by quite distinct underlying mechanisms.

Rather than being a magnesium mimic positioned to stabilize negative charge buildup on the β- and γ–phosphates at the catalytic transition state, direct visualization of the calcium binding-site by crystallography shows that calcium occupies a site near the α-phosphate (Figure 2B). In the closed state, the calcium is octahedrally coordinated by backbone carbonyl oxygens from the ATP-lid, Tyr221 hydroxyl and the ATP α-phosphate oxygen. Unlike the magnesium site, this site is configured quite differently in the apo state vs. the closed state, thus providing a unique coupling between open-closed conformation and calcium binding (Figure 3; Movie 1). TRAP1’s ability to close does not require divalent cations, since ATP alone is sufficient to stabilize the closed state and reach an identical asymmetric conformation^20,21^. Moreover, since dimer closure is rate-limiting in the ATPase cycle, any decrease in catalytic efficiency caused by the absence of a magnesium-stabilized transition state is not expected to dramatically alter the overall reaction rate. Whether there is an alternative hydrolysis pathway where solvent can stabilize the charge build up on the β- and γ-phosphates while the calcium activates a different attacking water is not clear. Structural data supports that the calcium-binding cavity near the α-phosphate presented here is conserved in the GHKL (<u>G</u>yrase, <u>H</u>sp90, histidine <u>K</u>inase and Mut<u>L</u>) ATPase superfamily. Hearnshaw et al. observed a potassium ion occupying the same pocket with a similar octahedral coordination in DNA gyrase^27^, and they pointed out that in yeast topoisomerase II, a lysine ε-ammonium group occupies the same pocket. For Hsp90s, perhaps this pocket is responsible for the mild ATPase stimulation by monovalent-cations previously observed for yeast Hsp90^28^.

**Figure 3.**
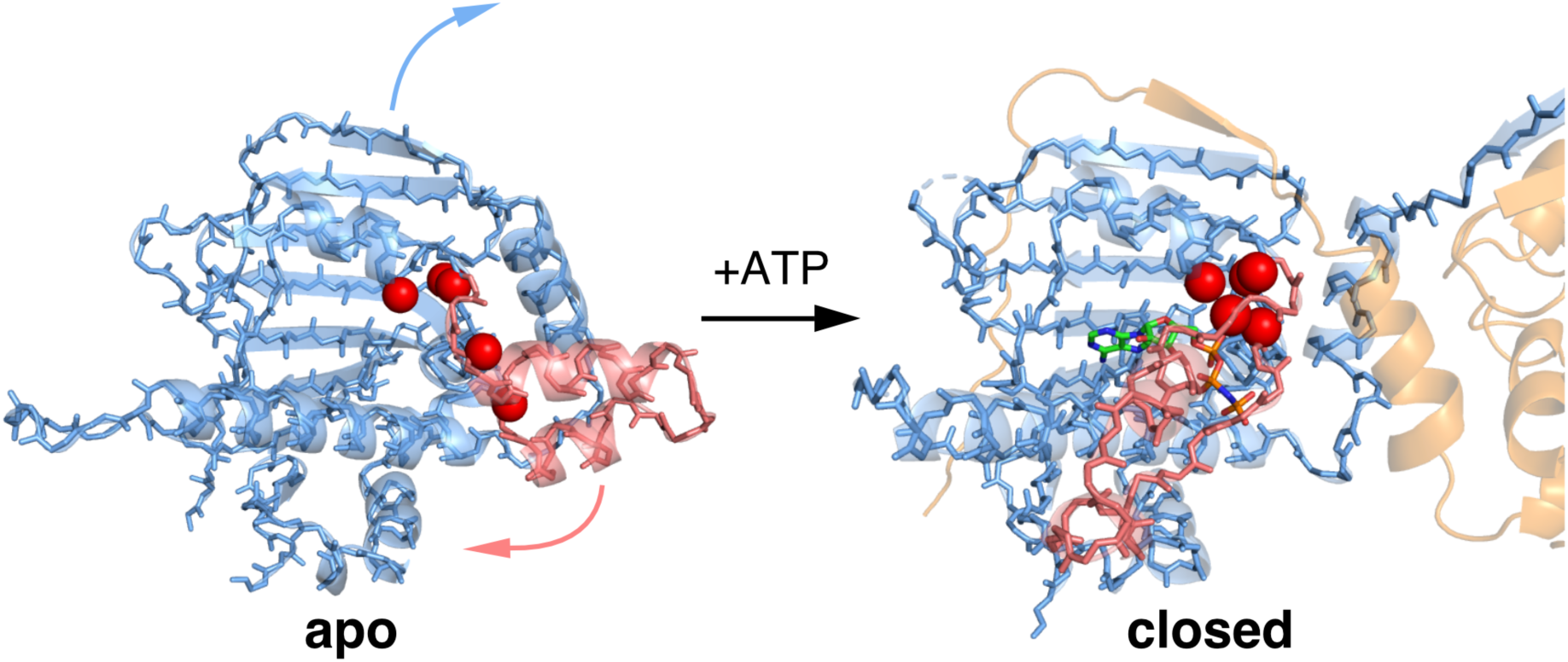
Molecular model for mechanism of calcium-binding in TRAP1. The lid region (pink highlight) adopts an open state in absence of nucleotide (apo, left). The oxygen atoms involved in calcium binding (red spheres) do not form a cluster in the apo state. Once ATP binds the NTD, closing of the ATP-lid rearranges these oxygen atoms and an α-phosphate oxygen from the ATP completes the calcium coordination in an octahedral geometry (closed, right). The movement of the lid (pink arrow) from open to closed is accompanied by the strand swap (blue arrow) that stabilizes interactions between NTDs (transparent blue and orange cartoon). In this model, the calcium-binding sites are only formed once ATP fully occupy both sites and the NTDs are dimerized, explaining the apparent cooperativity in calcium-binding.

The remarkable change in cooperativity observed with calcium indicates that TRAP1 can adopt a switch-like response to calcium at high ATP levels, providing a strong connection between calcium concentration in the mitochondrial matrix and TRAP1 ATPase-driven cycling. While the physiological consequences have yet to be determined, there are at least several scenarios where this could modulate cell function: i) by participating in the calcium-dependent switch between glycolysis and oxidative phosphorylation; ii) by modulating TRAP1’s role in opening the mitochondrial transition pore via altered sequestration of cyclophilin D binding, iii) by switching TRAP1 functionality or substrate specificity from client remodeling to more of a holdase behavior. Indeed, all of these could be a consequence of short-circuiting the normal, two-step ATP hydrolysis mechanism^20^, going directly from closure to full reopening, and skipping the flip in asymmetry believed to be coupled to client remodeling (Figure 4). TRAP1’s ability to suppress client aggregation by a capture and release mechanism (holdase activity) would likely be unaffected. Related to this, it would be interesting to see whether TRAP1 has a distinct subset of client proteins under high vs. low concentrations of ATP.

**Figure 4.**
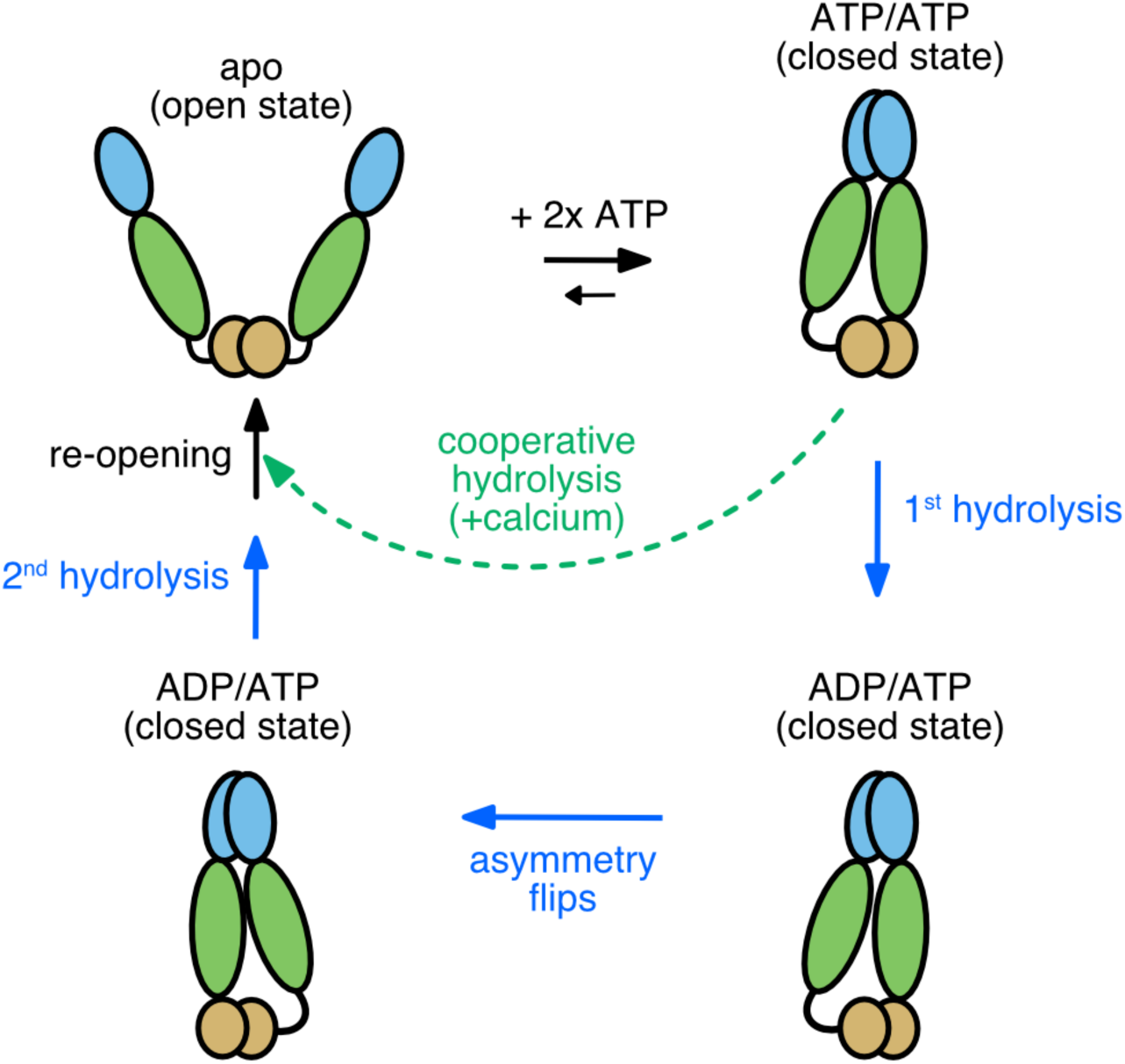
Model for TRAP1 ATPase cycle showing the normal sequential two-step ATP hydrolysis mechanism observed with magnesium (solid blue arrows and text) and an alternative path (dashed green arrow and text) when using calcium. The open dimer binds 2 ATPs, which in turn stabilizes an asymmetric closed state. Calcium-binding at least partially bypasses the magnesium-supported pathway for step-wise hydrolysis, resulting in cooperative ATPase activity that consequently skips the conformational switch that happens after the first ATP hydrolysis.

In principle, since calcium ions bind at an orthogonal site from magnesium, simultaneous binding of both divalent cations is possible. Preliminary experiments did not show synergistic effects when calcium is added to magnesium-ATPase reaction or vice-versa. While the physiological context for this calcium and magnesium activity modulation is not yet known, the findings in this work provide a molecular model to inform future work in elucidating the role of TRAP1 in the mitochondria.

## Materials and methods

### Protein purification

Full-length wild-type hTRAP1 and zTRAP1 (without mitochondrial signal sequence) was cloned into a pET151DTOPO vector as described previously^19^. Point mutations were introduced via site-directed mutagenesis by polymerase chain reaction. For FRET assays, single-cysteines (G151C and K428C) were introduced into a cysteine-free zTRAP1 construct to enable site-specific labeling with maleimide derivatives of AlexaFluor 555/647 (Life technologies). Proteins were expressed in BL21(DE3)-RIL *E. coli* cells grown in TB media by addition of 0.5 mM IPTG at OD600 ∼0.8, then incubated with shaking for 12-18 hours overnight at 16°C. All of the TRAP1 constructs have an N-terminal 6xHis-tag with a cleavable TEV-site. Purification follows a standard protocol for an Ni-NTA chromatography using 40 mM Tris buffer at pH 8.0, followed by an anion exchange with a MonoQ column, and a final size-exclusion step with a Superdex S200 16/60 (GE Healthcare) column.

### FRET assays

FRET measurements were taken on a FluoroMax-4 (HORIBA Jobin Yvon) using excitation wavelength 532 nm and emission wavelengths 567/668 nm for Alexa555/647, respectively. Temperature was maintained at 20°C by a circulating water bath connected to the sample chamber. Kinetics were recorded every 4 seconds with an integration time of 0.3 seconds. To initiate dimer closure, 10 mM ATP was added to the reaction. Hydrolysis was then triggered by addition of 10 mM MgCl_2_ or CaCl_2_.

### ATPase assays

Most of the ATPase assays performed in this study uses the phosphate release assay as previously described^20^ using a coumarin-labeled phosphate binding protein (PBP)^29^. In all assays, 0.5 μM of TRAP1 dimer was used per reaction. Heterodimeric hTRAP1 was formed by equilibrating a mixture of the wild-type protein with 15-fold molar excess of the D158N point mutant (the D158N homodimer is inactive). Fluorescence was measured using a SpectraMax M5 plate reader by exciting at 385 nm and measuring emission at 475 nm with a cutoff filter at 455 nm at ‘Low’ sensitivity setting. The ATPase assays with zebrafish TRAP1 also monitors phosphate release via a chromogenic substrate 7-methylthioguanosine (7-MESG) in presence of an *E. coli* PNPase (purine nucleoside phosphorylase). In these assays, 0.14 mM 7-MESG and 2 μM PNPase was used per reaction. The net absorbance of 7-MESG was measured by subtracting absorbance at 500 nm from 354 nm. Both the PBP and PNPase used in this study was expressed and purified as previously described^20^. Initial rates from phosphate release assays were from obtained from fitted slopes of a linear function, y = mx + b, to the most linear region of each kinetic trace. The activity vs. concentration curves were analyzed by a standard Michaelis-Menten equation with the Hill-coefficient, n, in the exponent:

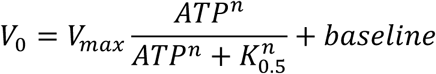

Where [ATP] is the ATP concentration and K_0.5_ is ATP concentration at which activity is half the maximum, V_max_. The two-population ATPase activity model of wild-type TRAP1 in magnesium used the following equation:

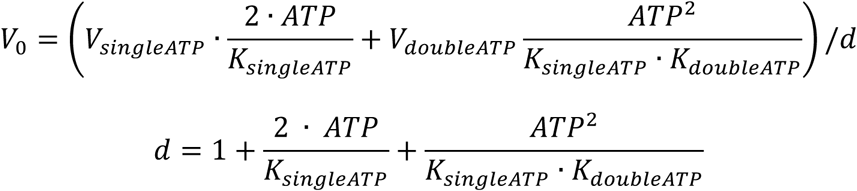

Where K_singleATP_ and K_doubleATP_ corresponds to apparent affinity constants at which half-maximum activity is achieved for half- (1-ATP bound) or fully-occupied (2-ATP bound) population, with V_singleATP_ and V_doubleATP_ is the corresponding estimated maximum activity.

### Crystallography and data processing

Zebrafish TRAP1 was incubated with 2 mM AMPPNP in presence of 2 mM CaCl_2_ at 5 mg/mL protein concentration. The reaction was incubated for at least 1 hour at 30°C to allow accumulation of the closed state prior to crystallization trials. Hanging drop reactions plate were set by mixing the protein with the crystallization solution (17% PEG-3350, 0.2 M Na/K tartrate, 36 mM hexammine cobalt) at 1:1 volume ratio (1 μL each) using a 15-well plate. Crystals grew in 2-3 days at room temperature. All diffraction data were collected at beamline 8.3.1 at the Advanced Light Source (ALS) in Berkeley, CA. Data collection and refinement statistics are in Table 4. Two wavelengths were used for the anomalous data collections: 11111 eV (1.116 Å) and 7450 eV (1.66 Å). Datasets were indexed using XDS^30^. The native dataset collected at shorter wavelength solved using molecular replacement of TRAP1 in the closed state^19^. Structures were refined using REFMAC5^31^ as part of the CCP4i package^32^. Anomalous difference map was generated using the program ANODE^26^ using input files generated by the program SHELXC^33^ and the solved structure without the calcium atoms.

**Table 4.** Data collection and refinement statistics for zebrafish TRAP1 bound to AMPPNP and calcium.

|  |  |
| --- | --- |
|  | **Zebrafish TRAP1 bound to AMPPNP and calcium** |
| **Wavelength** | 1.116 |
| **Resolution range** | 24.6 - 2.6 (2.693 - 2.6) |
| **Space group** | C 1 2 1 |
| **Unit cell** | 176.252 95.751 124.546 90 134.663 90 |
| **Total reflections** |  |
| **Unique reflections** | 44418 (4360) |
| **Multiplicity** |  |
| **Completeness (%)** | 97.71 (97.39) |
| **Mean I/sigma(I)** |  |
| **Wilson B-factor** | 59.92 |
| **R-merge** |  |
| **R-meas** |  |
| **R-pim** |  |
| **CC1/2** |  |
| **Reflections used in refinement** | 44407 (4358) |
| **Reflections used for R-free** | 2218 (183) |
| **R-work** | 0.1863 (0.3241) |
| **R-free** | 0.2366 (0.4081) |
| **Number of non-hydrogen atoms** | 9798 |
| **macromolecules** | 9557 |
| **ligands** | 64 |
| **solvent** | 177 |
| **Protein residues** | 1186 |
| **RMS(bonds)** | 0.016 |
| **RMS(angles)** | 1.31 |
| **Ramachandran favored (%)** | 94.28 |
| **Ramachandran allowed (%)** | 4.68 |
| **Ramachandran outliers (%)** | 1.04 |
| **Rotamer outliers (%)** | 9.26 |
| **Clashscore** | 10.05 |
| **Average B-factor** | 73.94 |
| **macromolecules** | 74.61 |
| **ligands** | 42.22 |
| **solvent** | 49.00 |
| **Number of TLS groups** | 7 |

## Acknowledgements

We thank members of the Agard Lab, especially Miguel Betegon for helpful discussions. We gratefully thank Rose Citron for initial help with crystal screening and data collection. Support for this work was provided by the NIH Protein Structure Initiative–Biology Grant U01 GM098254 and the Howard Hughes Medical Institute. We also thank James Holton, George Meigs, and staff at Advanced Light Source (ALS) beamline 8.3.1 for help with data collection. Beamline 8.3.1 at the ALS is operated by the University of California Office of the President, Multicampus Research Programs and Initiatives grant MR-15-328599 and Program for Breakthrough Biomedical Research, which is partially funded by the Sandler Foundation.

